# Active Site Heterogeneity Drives Differential Sialic Acid Metabolism in *Enterobacteriaceae* Pathogens

**DOI:** 10.1101/2025.09.25.678553

**Authors:** Sai Rupini Vemuri, Harish Babu Kolla, Sai Archana Kurucherlapati, Prakash Narayana Reddy

## Abstract

*Enterobacteriaceae* pathogens like *E. coli, Shigella* and *Salmonella* are mainly associated with the food poisoning and various mucosa associated infections. These pathogens invade the host and rely on the host derived nutrients for their survival. One such nutrient is sialic acid (N-acetylneuraminic acid is the common one) which is present in the mucosal linings throughout the body. Ability of these pathogens to breakdown and utilize sialic acid for their growth is a crucial factor for their survival and pathogenesis. However, the heterogeneity among these enteric pathogens remains unstudied. In this study, we explored the difference in the sialic acid utilization pattern among these pathogens and found that the interactions between the N-acetylneuraminic acid (Neu-5-Ac) and the NanA, an enzyme that catalyze the breakdown of sialic acid play a key role for this heterogeneity. Additionally, we found that the Neu-5-Ac interacted differently with the NanA from each strain because of the variations in the active sites/contact sites which made the strains to utilize sialic acid different. Finally, our results provide an insight into the current understanding of sialiobiology research by highlighting the active site/contact sites differences drived the pathogens’ ability to adapt to diverse host environments, potentially altering their pathogenicity and resistance to host immune responses. This study provides new insights into the evolution of *Enterobacteriaceae* and suggest potential targets for therapeutic intervention aimed at disrupting sialic acid utilization.

## Introduction

The *Enterobacteriaceae* pathogens like *Salmonella, Escherichia coli* O157:H7 and *Shigella* sp are known to thrive in the complex ecosystem of the human gastrointestinal tract^1,2^. Along with adapting to the host environment, these pathogens are characterised by their ability to exploit host-derived nutrients to sustain within the host and initiate infections. Sialic acid, a family of 9-carbon acidic sugars is one such nutrients proved to be consumed by the pathogens^3^. It is found to be located on the surface of the host cells, particularly the mucosal glycoconjugates. Sialic acid has been found to have a critical role in immune modulation, influencing host-pathogen interactions and pathogen virulence^4^. Besides, sialic acid is an important carbon source for the sustenance and the growth of the bacteria within the human host. Pathogens like the *Enterobacteriaceae* have evolved complex mechanisms to utilize the sialic acid.

On the host mucosal lining, the sialic acid is present as the terminal component on the glycoproteins and gangliosides, it is also found to be present in many secretions like mucous and saliva. The pathogens have grown to imbibe systems that enable them to obtain sialic acid by the action of sialidases (also known as Nan enzymes)^5^. These enzymes work by cleaving the sialic acid from the host glycoconjugates. The N-acetylneuraminic acid lyase (*NanA)* is involved in the initial step of the breakdown of the sialic acid and convert to N-acetylmannosamine by removing a pyruvate group^5^. This intermediate is later catabolized by a series of steps to produce fructose-6-phosphate to drive glycolysis and produce energy for the cell survival^5^.

Several studies revealed polymorphism in the sialic acid catabolism genes that lead to the differential utilization pattern of sialic acids^5,6^. However, the degree of sialic acid metabolism is unstudied in the most virulent pathogens of *Enterobacteriaceae* like *E. coli* O157: H7, *Shigella* sp, and *S. typhimurium*. The differential ability to uptake and metabolize sialic acid can influence virulence, immune evasion, and persistence within the host, with some pathogens exhibiting a more efficient capacity for sialic acid utilization than others. Understanding the molecular mechanisms that underlie this heterogeneity in sialic acid metabolism is crucial for developing targeted therapeutic strategies against *Enterobacteriaceae* infections.

## Materials and methods

### Bacterial culture, media, and reagents

The brain heart infusion (BHI) and M9 minimal media and N-acetylglucosamine were procured from Sigma Aldrich (Guntur distribution, Andhra Pradesh, India). The standard strains of *Enterobacteriaceae* used in this study were from our previous institute Vignan’s Foundation for Science, technology, and Research, Guntur, Andhra Pradesh, India during March 2022 where we used these strains in our previous studies **(Table 1)**. The bacterial cultures were propagated, and their glycerol stocks were maintained for the future use. Briefly, the cultures were streaked on the brain heart infusion (BHI) agar and inoculated in the BHI broth and incubated overnight at 37^°^C with constant agitation.

**Table 1.**
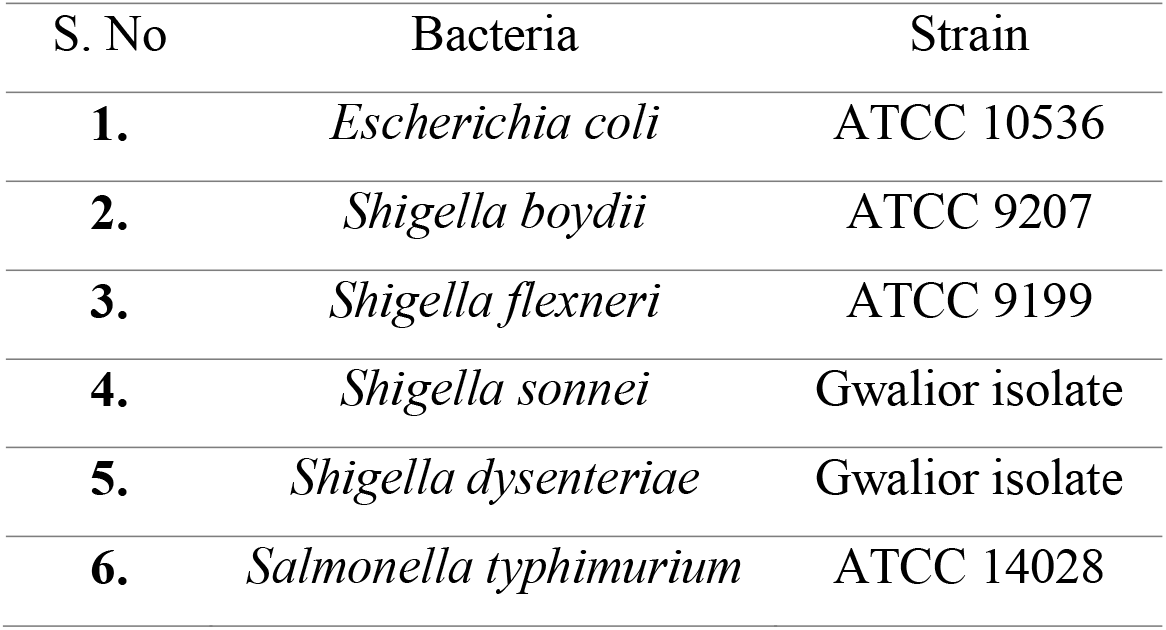
List of bacterial strains used in the study.

| S. No | Bacteria | Strain |
| --- | --- | --- |
| 1. | *Escherichia coli* | ATCC 10536 |
| 2. | *Shigella boydii* | ATCC 9207 |
| 3. | *Shigella flexneri* | ATCC 9199 |
| 4. | *Shigella sonnei* | Gwalior isolate |
| 5. | *Shigella dysenteriae* | Gwalior isolate |
| 6. | *Salmonella typhimurium* | ATCC 14028 |

### Bacterial culture

Around 100 microliters of overnight culture were inoculated in to 5 mL of M9 minimal media supplemented with 1 mg/mL of Neu-5-Ac and allowed to culture for 24 hrs in a 37^°^C shaking incubator^5^. After 24 hrs, the OD values of each culture in the presence of Neu-5-Ac was measured at 600 nm.

### Bioinformatics analysis

The complete amino acid sequences of NanA protein of the *E. coli O157:H7, Shigella sp and S. typhimurium* was obtained from Uniprot data base (https://www.uniprot.org/). These amino acid sequences were aligned to determine the degree of sequence homology and phylogenetic relationship by multiple sequence alignment using Clustal Omega online resource (https://www.ebi.ac.uk/jdispatcher/msa/clustalo). The Structural analysis was performed for the molecular docking studies. The three-dimensional structures of NanA protein were determined using Raptor-X server (http://raptorx.uchicago.edu/)^7^. Later, these structures were refined to the experimental quality through Galaxy-Refine2 server (http://galaxy.seoklab.org/refine2)^8^. The refined protein structures were then used for molecular docking studies.

Molecular docking was carried out to determine the two-dimensional interactions between the NanA protein and Neu5Ac ligand. For this, the three-dimensional structure of Neu5-Ac was retrieved from pubchem data base in SDF format. Docking analysis was carried out using CB-DOCK2 tool (https://cadd.labshare.cn/cb-dock2/index.php)^9^ and the binding affinity and 2-D interactions between each complex were noted.

## Results

### Bacterial growth curve

The OD values at 600 nm were recorded to determine the bacterial growth after 24 hrs of culture with Neu-5-Ac as a sole carbon source in the M9 minimal media. The OD values indicate that all the strains had utilized the Neu-5-Ac for their growth *in vitro* **(Figure 1)**. The bar graphs shown in figure 4 indicate that the *E. coli O157:H7* and *S. dysenteriae* has shown maximum growth in the presence of Neu-5-Ac as compared to the others **(Figure 1)**. This finding signifies that these strains as the potential sialic acid utilizers among the considered *Enterobacteriaceae* members while *S. typhimurium* and *S. boydii* in the next place **(Figure 1)**. The other two species of *Shigella-sonnei* and *flexneri* has shown the same range of growth with the Neu-5-Ac **(Figure 1)**. However, these readings are significant as compared to the control with M9 media alone. Overall, the *in vitro* growth curve data shows that the *Enterobacteriaceae* strains used in this study has potentially utilized Neu-5-Ac as a carbon source for their growth.

**Figure 1.**
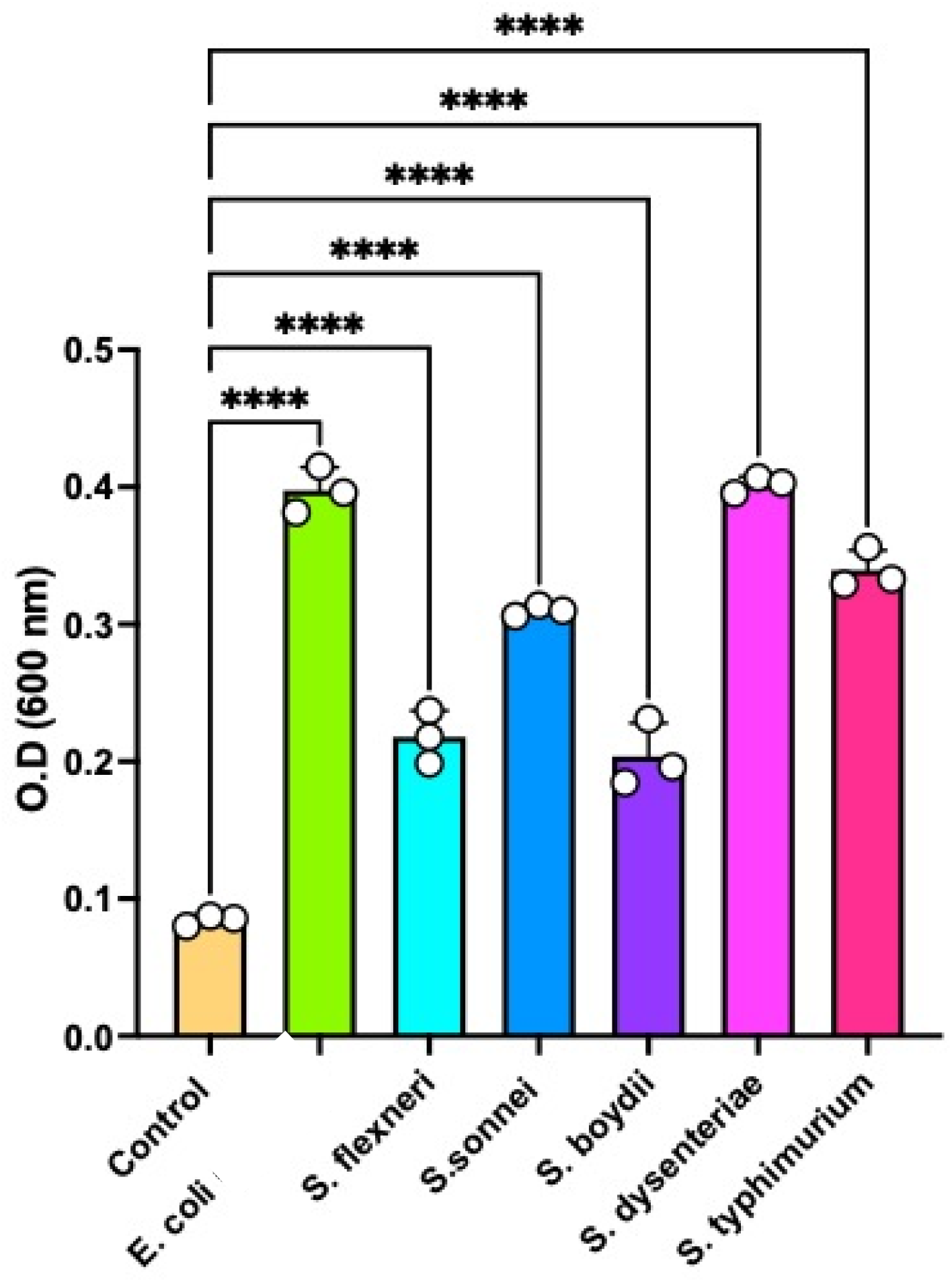
OD values showing the bacterial growth in the M9-minimal media after 24 hrs of culture with Neu-5-Ac.

### Sequence homology and phylogeny

Based on the bacterial growth curve, it is appreciable that the strains have shown difference in the Neu-5-Ac utilization pattern after 24 hrs. Now, the next question we asked is whether there is any genetic or molecular mechanism that underlies this observation. For this, we have performed in silico analysis based on the genome sequence information available in the online databases. One such is Uniprot, where we obtained the complete amino acid sequence of the NanA protein. Here we tried to see if there are any variations in the NanA protein among these *Enterobacteriaceae* strains that could be a reason for the differential utilization pattern of Neu-5-Ac. Multiple sequence alignment of NanA protein revealed it as a highly conserved protein with a significant variation **(Figure 2) (Table 2)**. Almost 90 % of the amino acids are conserved in the NanA protein among these strains with variations throughout the protein.

**Table 2.** Amino acid sequence identity in the NanA protein among the *Enterobacteriaceae* members.

|  | *S. typhimurium* | *S. boydii* | *E. coli* O157:H7 | *S. flexneri* | *S. sonnei* | *S. dysenteriae* |
| --- | --- | --- | --- | --- | --- | --- |
| *S. typhimurium* | 100.00 | 90.24 | 90.24 | 90.24 | 90.57 | 90.57 |
| *S. boydii* | 90.24 | 100.00 | 99.33 | 99.33 | 99.66 | 99.66 |
| *E. coli* O157:H7 | 90.24 | 99.33 | 100.00 | 99.33 | 99.66 | 99.66 |
| *S. flexneri* | 90.24 | 99.33 | 99.33 | 100.00 | 99.66 | 99.66 |
| *S. sonnei* | 90.57 | 99.66 | 99.66 | 99.66 | 100.00 | 100.00 |
| *S. dysenteriae* | 90.57 | 99.66 | 99.66 | 99.66 | 100.00 | 100.00 |

**Figure 2.**
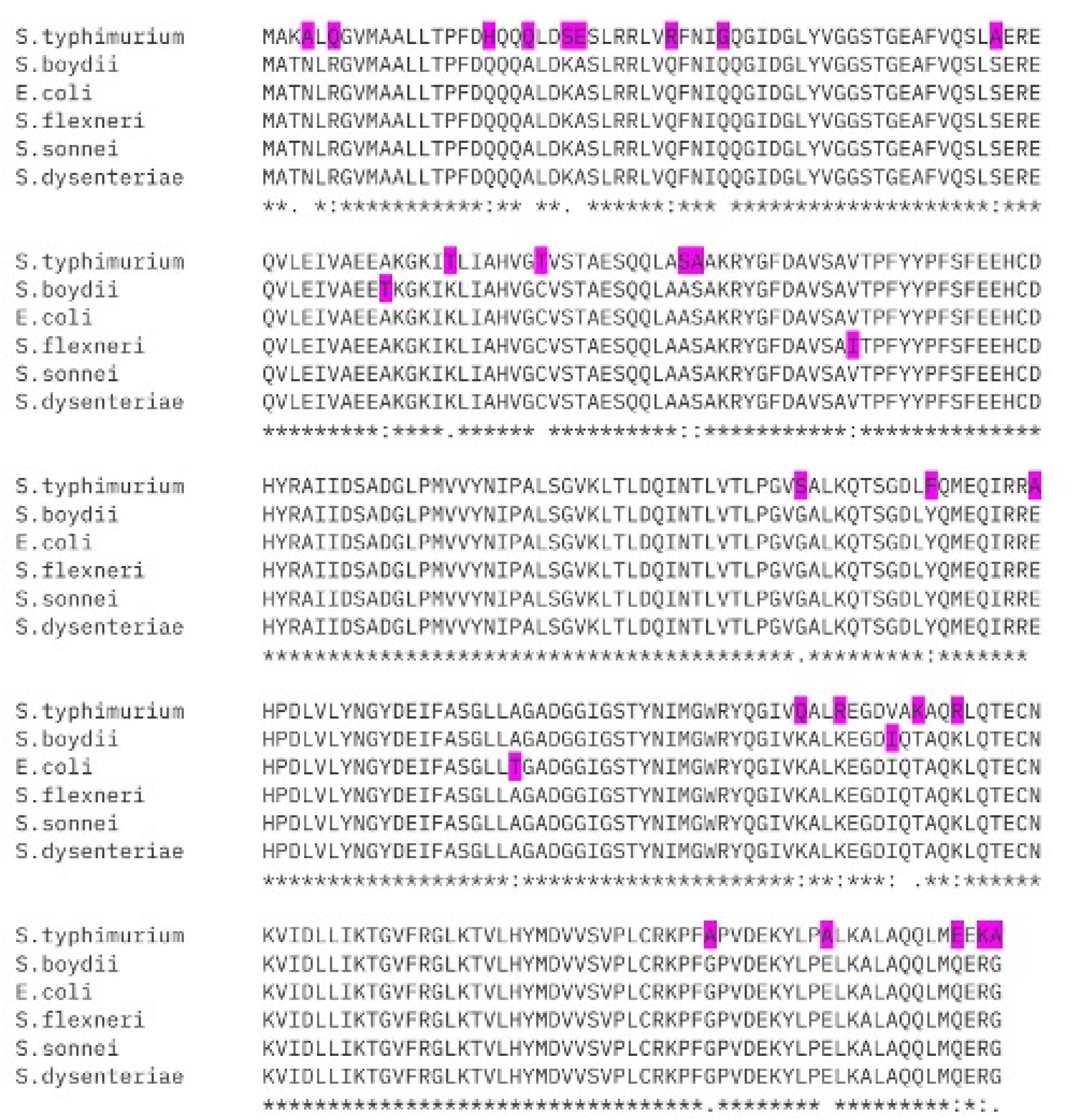
Multiple sequence alignment of NanA sequences of *Enterobacteriaceae* members.

On the other hand, Phylogenetic analysis revealed that the NanA is very closely related among *Shigella* sp and *E. coli* O157:H7 while *S. typhimurium* in the monophyletic group in the tree **(Figure 3)**. This finding is aligning with several previous reports on the comparative genetic analysis of *E. coli* and *Shigella* sp^10-12^.

**Figure 3.**
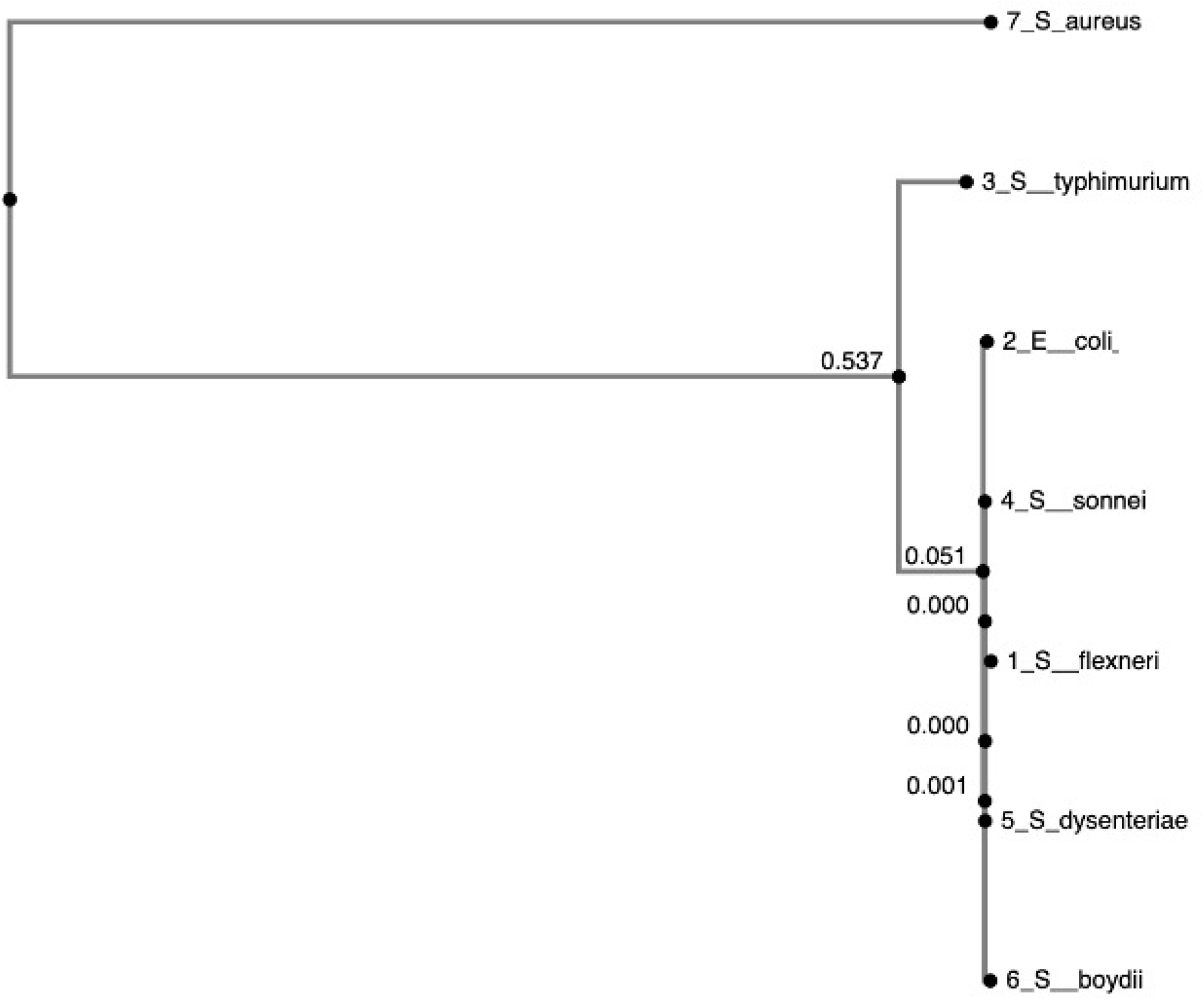
Phylogenetic tree of NanA protein of *Enterobacteriaceae* members.

### Molecular docking analysis

As the sequence and phylogenetic analysis revealed the variations in NanA protein among the *Enterobacteriaceae* strains, we later investigated whether these variations are at the active sites that would bring differences in the catalysis, breakdown and the final growth of the bacteria. For this we used molecular docking as a potential tool to identify the contact/active sites in the NanA protein and the interaction pattern with Neu-5-Ac.

Molecular docking was carried out by predicting the three-dimensional structure of NanA protein followed by refinement. The refined structures were docked with the Neu-5-Ac ligand and the findings were recorded. The binding affinities of Neu-5-Ac with the NanA of *E. coli* O157:H7, *S. flexneri, S. dysenteriae, S. boydii, S. sonnei* and *S. typhimurium* are as follows: −5.5, −5.8, −5.8, −4.6, −6.8, and −6.3 Kcal/mol respectively with the interacting amino acids as *E. coli* O157:H7: ALA11 SER47 THR48 ILE139 LEU142 LYS165 THR167 GLY189 TYR190 ASP191 GLU192 ILE206 GLY207 SER208 THR209 ILE243 ILE247 VAL251, S. flexneri: ALA11 GLY46 SER47 THR48 GLU50 ALA51 PHE52 VAL53 PHE109 TYR110 TYR137 ILE139 ALA141 LEU142 LYS165 THR167 GLY189 TYR190 ASP191 GLU192 ILE206 GLY207 SER208 THR209 ILE243 ILE247 VAL251 PHE252 LEU255 PRO273 PHE274, S. dysenteriae: ALA11 THR48 TYR137 ILE139 LYS165 THR167 GLY189 TYR190 ASP191 GLU192 ILE206 GLY207 SER208 THR209 ILE243 ILE247 VAL251 LEU255, S. boydii: ALA11 SER47 THR48 TYR137 ILE139 LEU142 LYS165 THR167 GLY189 TYR190 ASP191 GLU192 ILE206 GLY207 SER208 ILE243 ILE247 VAL251 PHE252 LEU255, S. sonnei: ALA11 GLY46 SER47 THR48 PHE109 TYR137 ILE139 ALA141 LEU142 LYS165 THR167 GLY189 TYR190 ASP191 GLU192 ILE206 GLY207 SER208 THR209 ILE243 ILE247 VAL251 PHE252 LEU255, and S. typhimurium: ALA11 SER47 THR48 TYR137 ILE139 ALA141 LEU142 LYS165 THR167 SER168 GLY169 GLY189 TYR190 ASP191 GLU192 ILE206 GLY207 SER208 THR209 ILE243 ILE247 VAL251 PHE252 LEU255 PHE274 as shown in **table 3**.

**Table 3.**
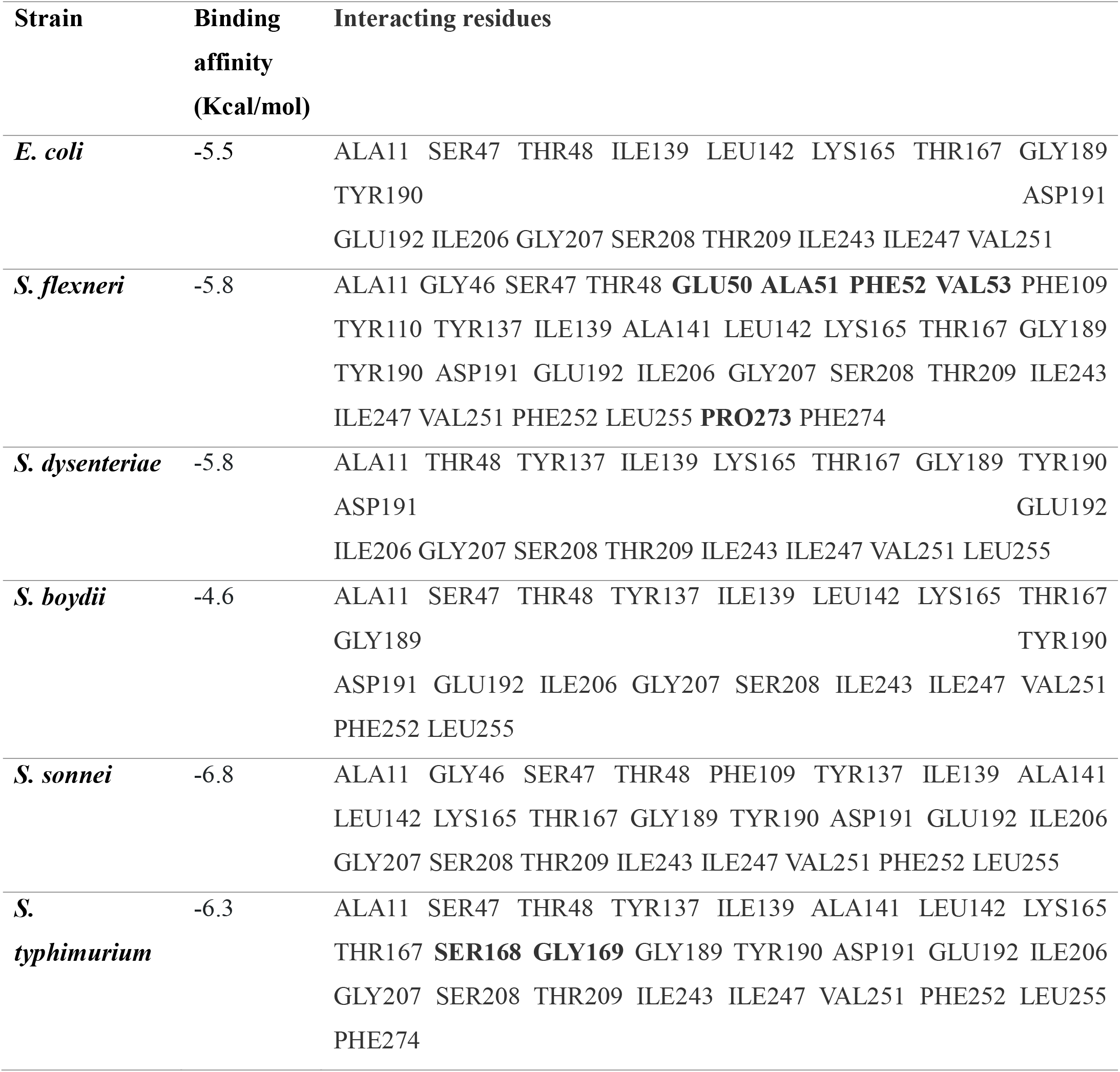
Binding affinities and the amino acid residues in NanA protein interacting with the Neu5Ac.

These findings indicate that the Neu-5-Ac binds differently to the NanA of *E. coli* O157:H7, *S. flexneri, S. dysenteriae, S. boydii, S. sonnei* and *S. typhimurium* based on their active sites and hence the difference in the binding affinities which lead to the differential utilization pattern of Neu-5-Ac by these bacteria **(Figure 4)**.

**Figure 4.**
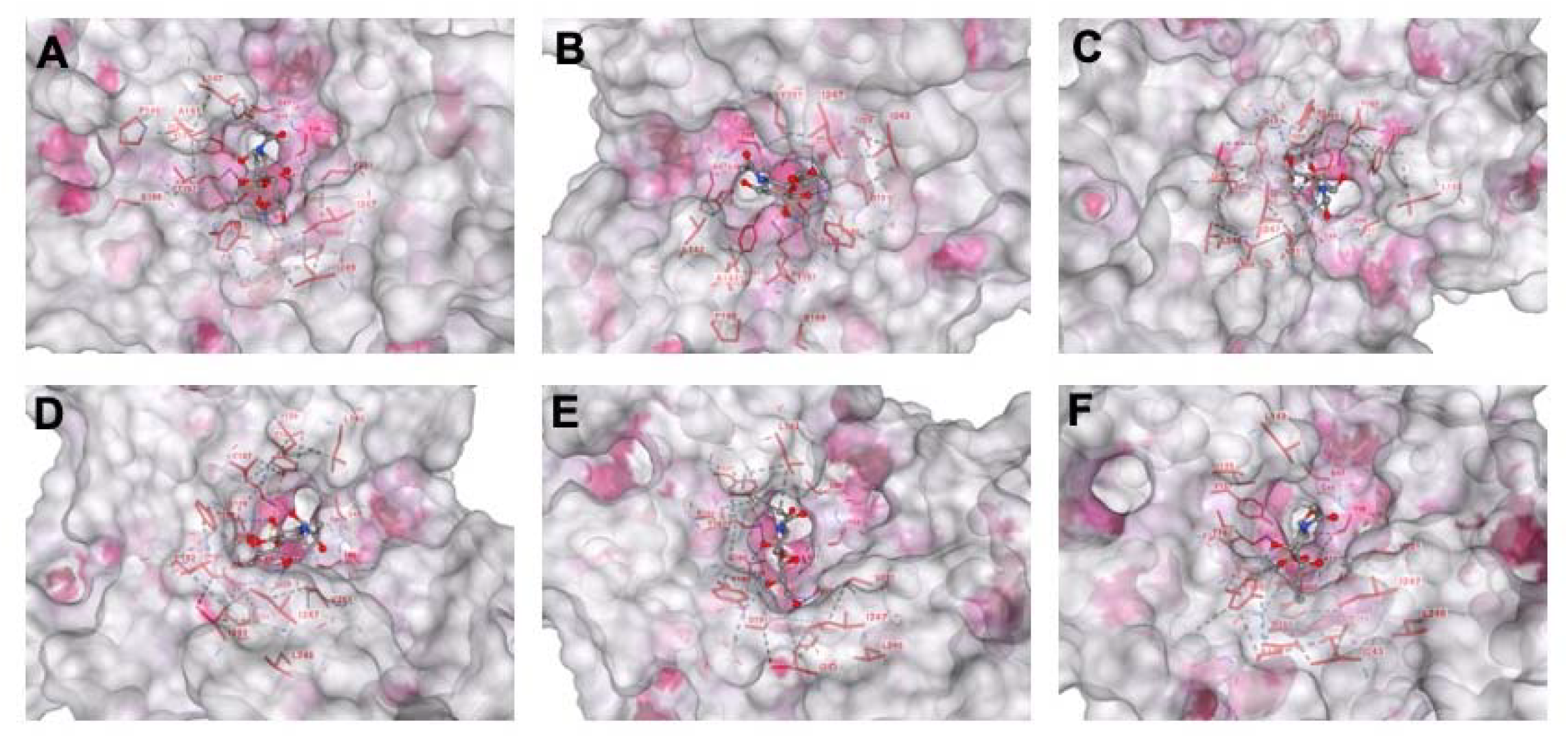
Molecular docking analysis showing the interaction between Neu-5-Ac and NanA of **(A)** *E. coli*, **(B)** *S. flexneri*, **(C)** *S. dysenteriae*, **(D)** *S. boydii*, **(E)** *S. sonnei*, and **(F)** *S. typhimurium*.

## Discussion

Although many studies on sialic acid role in pathogens are present, the effect of the enzyme *NanA* sialidases in *Enterobacteriaceae* pathogens has not been completely investigated. The presence of different amino acid sequences variations in the *NanA* protein in the *Enterobacteriaceae* pathogens resulting in the differential sialic acid utilization *in vitro* has been hypothesised. The amino acid sequence variations lead to the alteration in the factors such as the catalytic efficiency, substrate specificity or regulation resulting in different sialic acid catabolism mechanisms in different pathogenic strains. This diversity in the utilization in sialic acid suggest the effect on the ability to compete for sialic acid in the host environment, their total virulence potential and influence their mechanisms with the host cells^13^.

This study aims to investigate the various patterns in sialic acid utilization with *Salmonella, E. coli* O15:H7, and *Shigella* spp, particularly on the amino acid variation on the *NanA* protein. The catalytic sites of *NanA* proteins and bacterial growth from different strains of the pathogens are compared to find out the mechanism of specific mutations in the *NanA* protein give rise to the difference in the utilization of sialic acid *in vitro*.

The sequence variation exhibited by the *NanA* throughout the different strains of *Enterobacteriaceae* leads to the observed different pathways in their ability to use sialic acid. The genetic variability results in the functional change by modifying the amino acid sequence in the active site of the protein structural configuration which further impacts the degree of enzyme affinity for sialic acid and its catalytic efficiency. The structural diversity in NanA sialidases across *Salmonella, Shigella*, and *E. coli* O157:H7 is likely a major determinant of their sialic acid utilization profiles. Our comparative analysis revealed that the amino acid differences in NanA among these pathogens lead to distinct enzymatic activities, particularly in the hydrolysis of Neu5Ac. These differences likely reflect the unique evolutionary adaptations of each pathogen to their respective ecological niches, such as the intestinal environment or host-specific immune responses.

The diverse patterns of utilizing the sialic acid observed in these pathogens greatly impact other factors like pathogen colonization, persistence and host virulence. Furthermore, the differential binding affinities observed between the sialidases of these pathogens and Neu5Ac suggest that specific adaptations to host glycosylation patterns may govern the pathogen’s ability to utilize sialic acid. The amino acid interactions within NanA play a pivotal role in determining enzyme-substrate binding affinities and, consequently, the efficiency of sialic acid degradation. Our data suggests that these interactions not only affect the catalytic turnover of Neu5Ac but may also modulate the structural integrity of the enzyme, impacting its stability and function in different environmental conditions. These findings are consistent with previous studies highlighting the importance of NanA’s active site architecture and the presence of conserved motifs that facilitate substrate recognition.

Other than utilizing sialic acid for host survival, the pathogens also find it crucial for evading the host immune responses^4,14,15^. When the sialic acid component is removed from the host glycoproteins, the *Enterobacteriaceae* pathogens can change the glycan composition on their surface structures like lipopolysaccharides and outer membrane proteins. Through this mechanism, the pathogens can imitate the host cell surface, preventing any host immune cells recognition. Moreover, a free sialic acid, generated by the action of *NanA* sialidases is used as a signaling molecule involved in many survival processes like biofilm formation, modulation of inflammatory response and host immune evasion. Therefore, understanding the genetic variations and the consequent mechanisms in *Enterobacteriaceae* pathogens is very significant with regard to the basic microbiology and also aiding to develop therapeutic strategies to disrupt the sialic acid utilization pathways and inhibiting the virulence of these human pathogens is the final aim of our findings.

## Conclusion

Our study provides valuable insights into the molecular mechanisms underlying the utilization of sialic acid by *Enterobacteriaceae* pathogens. The variations in NanA structure and function among *Salmonella, Shigella*, and *E. coli* O157:H7 reflect the pathogen-specific adaptations that contribute to their virulence and survival in host environments. Understanding these differences at the molecular level will be crucial for developing targeted strategies to combat infections caused by these pathogens and to further elucidate the role of sialic acid in host-pathogen interactions.

## Acknowledgments

The authors are thankful for the management of Vignan’s Foundation for Science, Technology, and Research for providing with the cultures to pursue this study. PNR is thankful to Dr. V. Saleem Basha, Principal, SKR & SKR Govt. Degree College for Women (A), Kadapa, for his support and encouragement.

## Conflict of interest

The authors declare that there is no conflict of interest.

## Author’s contribution

SRV and HBK performed the experiments with the guidance of PNR. PNR conceived the study, planned experiments. SRV and SAK wrote and revised the paper. All the authors have approved the paper for the publication.

## Funding

Not applicable.

## Data availability

All datasets generated or analyzed during this study are included in the manuscript.

## Ethics statement

Not applicable.

## Notes

### Competing Interest Statement

The authors have declared no competing interest.

